# Interleukin 12 correlates with performance, metabolism and acid-base balance during physical exercise

**DOI:** 10.1101/2023.02.15.528787

**Authors:** Ione Vieira Castilho, Luis Carlos Oliveira Gonçalves, Shirley Gomes Leite Rodolpho, Jaqueline Santos Silva Lopes, Eduardo Luzia França, Adenilda Honório França, Aníbal Monteiro de Magalhães Neto

## Abstract

Studies involving physical exercise are no longer performed only to evaluate the performance of athletes, but have become an important tool to understand how different forms of stress affect immunometabolism. The present study investigated the acute impact of a treadmill running test on different biomarkers, the acid-base system, glycemia/lactatemia, and the correlation between IL-12 and metabolism/performance. Ten male subjects participated in a cross-sectional study. The treadmill protocol was progressively increased until exhaustion. The IL-12 concentration was measured using the “Cytometric Bead Array” kit (CBA, BD Bioscience, USA) through flow cytometry, and the data were analyzed using FCAP Array software. The test had an average time of 13 minutes and 51 seconds and induced alterations in IL-12 concentration of 160%, lactate of 607%, blood glucose of 58%, blood pH of −3%, BE of −529%, bicarbonate of - 58%, and anion gap of 232%. It was observed that the lower the percentage variation in IL-12, the greater the phase to reach the anaerobic threshold (AT) in Km/h, and the time to reach this same threshold, and the opposite was also true, confirmed by the Spearman test. (−0.900 between IL-12 and the time to reach AT and −0.872 between IL-12 and the phase to reach AT). Other correlations were observed: between post-IL-12 and pre anion gap of 1.0, post-IL-12 and post chloride of 1.0, percentage change in IL-12 and post anion gap of 1.0 and percentage variation in IL-12 and post lactate of 0.943, pre-IL-12 and post anion gap of −1.0, post-IL-12 and pre LDH of −0.943, post-IL-12 and post LDH of −0.943, post-IL −12 and BE post of −9.943, post-IL-12 and post bicarbonate of −0.943, and post-IL-12 and post pH of −0.943. The AT was reached in 7:52 minutes, in the 14.9 km/h phase, with a heart rate of 163 beats per minute, an absolute power of 524 W, and an absolute VO2 of 3.12 l.min. A correlation between IL-12 and performance, metabolism, and blood acid-base balance is suggested. Furthermore, it is expected that approximately 15% of glycemia is formed by the CORI cycle, through the removal of lactate and reestablishment of glycemia, however, this estimate can be exceeded in athletes, according to the level of training.

## Introduction

It has been some time since the models of study involving physical exercise ceased to be applied only to assess the performance of athletes, mainly with the emergence of two new fields of research, sportomic and immunometabolism, which have become important tools to understand how different forms of stress affect metabolism and the immune system and possible adaptations [1–5].

The sportomic method aims to generate real competition situations during an experiment, with the objective of understanding the metabolic and immunological alterations, and stress in different organs and systems, as well as selecting the most sensitive biomarkers for each sex, age, training level, type of stress, environmental conditions, and types of exposure [1,6,7,8,9,10].

Immunometabolism seeks to understand how the immune system and metabolism are affected by different situations, the acute and chronic responses, as well as how each modulates the other within a complex physiological system, such as the human body [11–14].

The different forms of integration between metabolic pathways, cells, tissues, systems, and biomarkers have intrigued researchers over time, still representing a current and innovative scientific field [15–19].

Among the main modulators, cytokines stand out, involved in almost all biological processes, including embryonic development, disease pathogenesis, response to infection, cognitive functions, progression of degenerative aging processes, stem cell differentiation, and the effectiveness of vaccines [20].

Among the research fields involving cytokines, one line has aroused the interest of many research centers around the world; the relationship between these modulating agents and physical exercise [21]. However, very little is known about the subject.

Among the known cytokines, interleukin-12 (IL-12) is a pro-inflammatory cytokine that regulates the responses of T cells and natural killer cells, induces the production of interferon-γ (IFN-γ), favors cell differentiation of T helper 1 (TH1) cells, and is an important link between innate resistance and adaptive immunity [22]. Considering research into the triad of exercise, IL-12, and immunometabolism, science is still taking its first steps [23,24].

Thus, the present study investigated the acute impact caused by a treadmill running test, on different biomarkers, changes in the acid-base system, and glycemia/lactatemia, as well as a possible relationship between IL-12 and metabolism/performance in running athletes.

## Material & methods

### Study Participants

The study included 10 male individuals, who are high-performance athletics athletes, without recent musculoskeletal injuries or any other health conditions that represent medical contraindications for the practice of exercise, and who consistently participate in municipal, state, and national competitions and have won medals. Anthropometric characteristics are presented in table 1.

**Table 1.** Characteristics of the participants (n = 10).

|  | Mean | Median | SD | SE |
| --- | --- | --- | --- | --- |
| **Age (years)** | 21.2 | 19.5 | 5.5 | 1.7 |
| **Weight (kg)** | 60.6 | 59.0 | 6.4 | 2.0 |
| **Stature (cm)** | 173.5 | 171.0 | 7.5 | 2.4 |
| **BMI (kg/m<sup>2</sup>)** | 20.1 | 20.1 | 1.4 | 0.4 |

**BMI** - Body mass index; **SD** - Standard Deviation; **SE** - Standard error; **n** – number of participants.

### Ethical Aspects

The participants received all the information about the objectives, procedures, and risks of the study and after agreeing, they were invited to sign an informed consent form (TCLE), which ensured their rights of privacy and freedom to withdraw from the study at any time they deemed necessary. The study was previously submitted to and approved by the Ethics Committee in Research involving Human Beings of the Federal University of Mato Grosso (UFMT), Araguaia campus, under opinion number: 2,230,073.

### Study Design

This is a cross-sectional study. Data collection was performed at the Cardiology Institute; the cytology analyses were carried out in the Chrono-immunomodulation laboratory, at the Federal University of Mato Grosso (UFMT), Araguaia campus; and the other blood analyses were carried out in a private laboratory, all located in the city of Barra do Garças – MT. Collections were performed between February and April 2021, always taking place at a standardized time, between 2:00 pm and 3:30 pm, in order to avoid possible influences of the circadian cycle. In addition, the ambient conditions were standardized, with the temperature maintained at 21° degrees.

All participants were instructed not to perform intense exercise in the 24-hour interval prior to the collection day and to eat a light meal two hours before the procedures. Each participant attended the collection site only once, and all the necessary procedures were carried out during that visit.

### Description of the Procedures

#### Exercise Protocol

As a basic warm-up, the athletes performed dynamic and static stretching in all body segments, lasting 5 minutes.

The treadmill exercise protocol was progressively increased until exhaustion, using a gas analyzer. The specific warm-up that preceded the test consisted of light running, which lasted 15 minutes. Afterwards, the athletes ran for an average total distance of 3 km, which was characterized by gradual load increases in stages, until voluntary exhaustion, with an initial speed of 10 km/h and an increment of 1 km every 400 meters.

In turn, the anaerobic threshold was determined by ventilatory parameters. The parameters used from the applied exercise protocol aimed to expose the participants to a distance similar to those faced in real field environments and trigger physiological alterations similar to those that would occur in a real running scenario.

#### Ergospirometric Evaluation

The cardiovascular assessment during the stress test was performed using an exercise electrocardiogram (ErgoPC Elite) with monitoring at lead points (D_1_, D_2_, D_3_, AV_R_, AV_L_, AV_F_, MC_5_, V_1_, V_2_, V_3_, V_4_, V_5,_ and V_6_). In addition, blood pressure (BP) was measured indirectly using a Missouri sphygmomanometer and a Rappaport stethoscope.

The metabolic evaluation was performed using a computerized analyzer (MetaLyzer 3B), a face mask (V2 mask Small), and software to capture and display the data, as well as to store and process all cardiorespiratory and metabolic variables evaluated.

It is important to mention that the evaluation equipment used was always calibrated between one participant and the next. The participants were positioned on the treadmill, wearing a face mask with a sterilized mouthpiece and the nose covered, with a specific fastener.

#### Flow Cytometry

The concentration of cytokine IL-12 present in serum samples was evaluated using the “Cytometric Bead Array” kit (CBA, BD Bioscience, USA). The analyses of these cytokines were performed using flow cytometry and the data analyzed using FCAP Array software.

#### Blood Samples

Two extractions of 5 mL each of venous blood were collected from the antecubital vein of each participant, by a qualified professional with experience in field collections. Blood collection took place with a stainless steel needle syringe, the first sample being extracted before the stress test and the second after the test.

After collection, the samples were stored in a metal-free polypropylene tube and subsequently centrifuged at 2500 rpm for 10 minutes at room temperature.

#### Statistical Analysis

Initially, descriptive statistics were performed on the data, with measurements of position (mean, median, mode, and percentiles) and dispersion (amplitude, variance, standard deviation, and standard error).

Afterward, the univariate analysis of these data was performed using the Shapiro-Wilk normality test (because the sample was smaller than 30 individuals). The equal variance test would be applied if the Shapiro-Wilk test presented a result indicating normal distribution (P>0.05). For results with P>0.05, the paired T-Student test would follow; if P≤0.05, the paired T-Student test would follow the non-parametric Mann-Witney test. If the Shapiro-Wilk test presented a result indicating non-normal distribution (P≤0.05), the non-parametric Mann-Witney test would be applied directly.

Still, in the phase of the univariate analysis, the analysis of repeated measures ANOVA One Way dependent was performed because they were the same individuals in different conditions and moments.

So, for a better interpretation of the data, the individuals were divided into two groups according to their sex. Then, the calculation of percentage variation was applied:

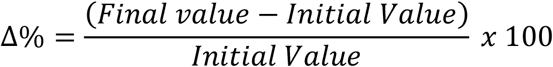

Cohen’s equations [25] were used to calculate the effect size for all variables to obtain Cohen d and r values:

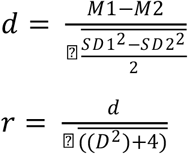

Where, M represent the means of observations and SD their respective standard deviations.

**Table 2.**
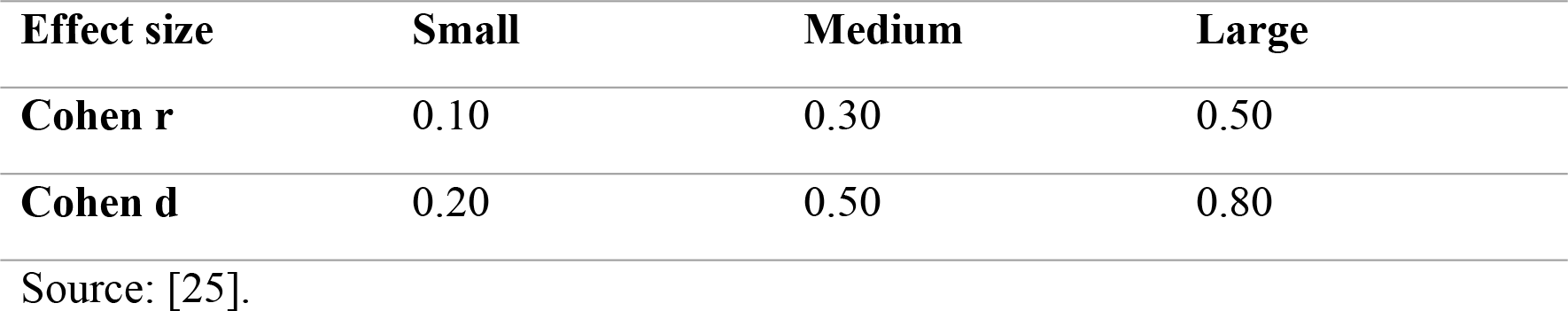
Values of effect size.

Next, multivariate data analysis was performed using data mining and machine learning techniques.

In this phase, in order to seek a bivariate measure between the data, because the observations contain quantitative values, the Pearson and Spearman correlation tests were applied, with the Spearman correlation being used for a visual analysis using the heat map strategy and the Pearson test as an initial measure for the following machine learning analyses.

As exploratory models of machine learning: CLUSTER - Classical Clustering (Agglomerative Hierarchical Method) and Nearest neighbor (single linkage); ORDINATION – Principal component Analysis (PCA) and Correspondence Analysis (CA).

The Z score was previously applied to adjust observations measurement units, and the Fruchterman-Reingold algorithm was applied with Euclidian Similarity Index [26,27].

SigmaPlot 14.5 (Academic Perpetual License - Single User – ESD Systat® USA) and, Past 4.03 (Free version for Windows) were used to carry out the different statistical tests and produce the graphs. Finally, the correlation coefficients were presented using heat maps [10].

## Results

The results referring to the exercise test and the cardiovascular aspects measured in the athletes that make up the sample are presented in supplementary table 1.

Supplementary table 2 presents the mean and standard error values for the pre and post exercise times, as well as the percentage variation between the means for the mentioned times. As the hematocrit showed a small percentage variation (4%) and did not present a significant difference between the means, it was not necessary to apply a correction factor to the values of all blood biomarkers, since when there is an acute change in the hematocrit, this indicates alteration in the state of hydration, with hemoconcentration or hemodilution, that would change the measured values, not due to a difference in its production, but due to a change in the amount of plasma.

Regarding the acid-base and hydroelectrolytic balance, a negative percentage variation for base excess (BE) of −529% is noteworthy, with a consumption of almost half of the bicarbonate and an increase in the anion gap of around 232%.

The treadmill test, with an average execution time of 13 minutes and 51 seconds, induced an increase in the concentration of interleukin 12 in the order of 160% and an increase in the concentration of lactate of 607%, revealing that despite the short performance time of the exercise, and the inclusion of high level athletes, the exercise can be considered as being high intensity.

Figure 1 shows the changes in glycemia and lactatemia induced by the test, with a percentage variation of 58% in glycemia and 607% in lactate, both with a significant difference, and a value of P<0.001.

**Figure 1.**
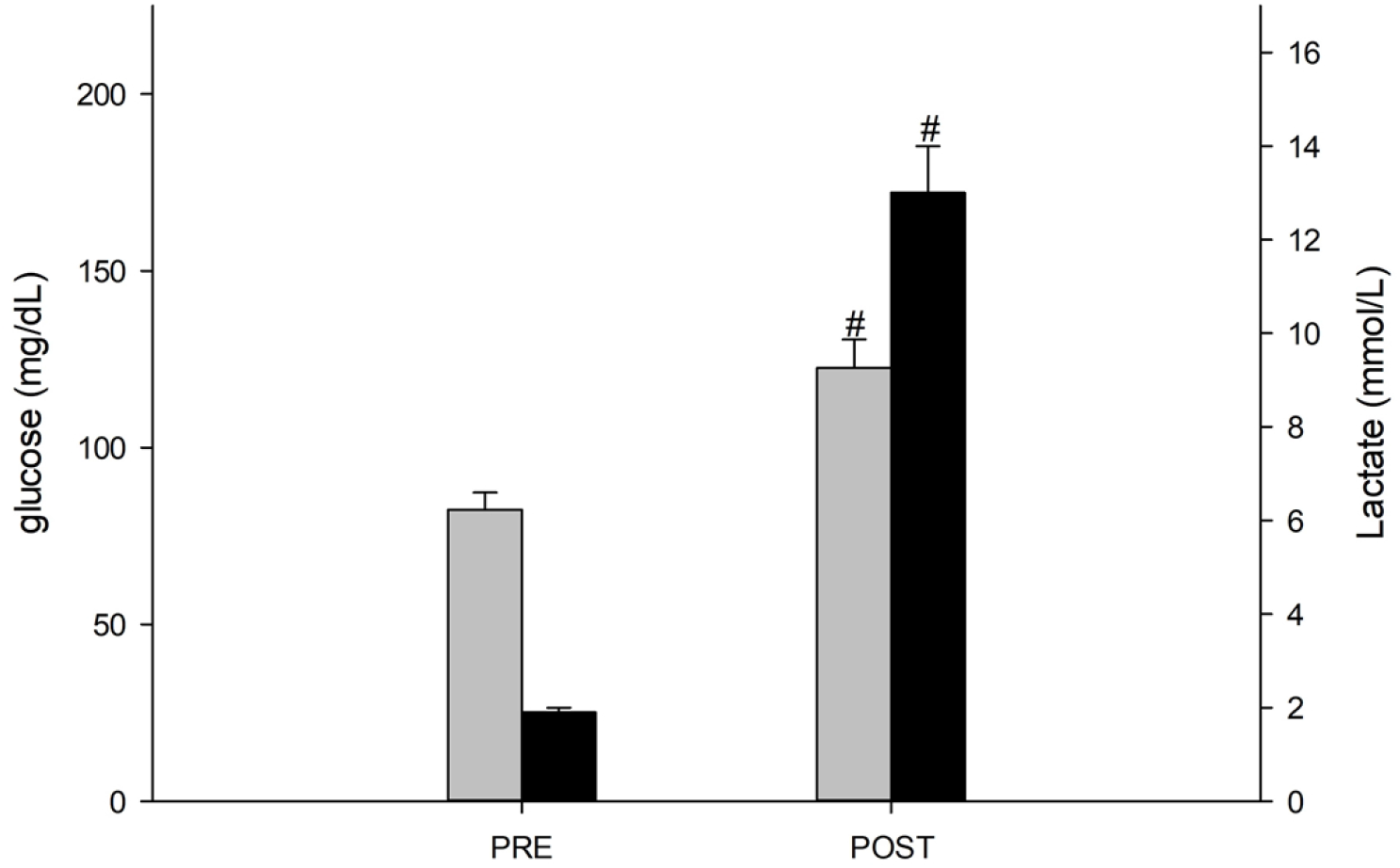
Protocol-induced alterations in blood glucose and lactate levels.

Regarding the acid-base balance, figure 2 summarizes the data presented in supplementary table 2, showing that the blood pH presented a small percentage variation (−3%), but with a significant difference (P<0.001), that the base excess showed robust variation (−529%), with a significant difference (P<0.001), and bicarbonate showed considerable variation (−58%), also with a statistical difference (P<0.001).

**Figure 2.**
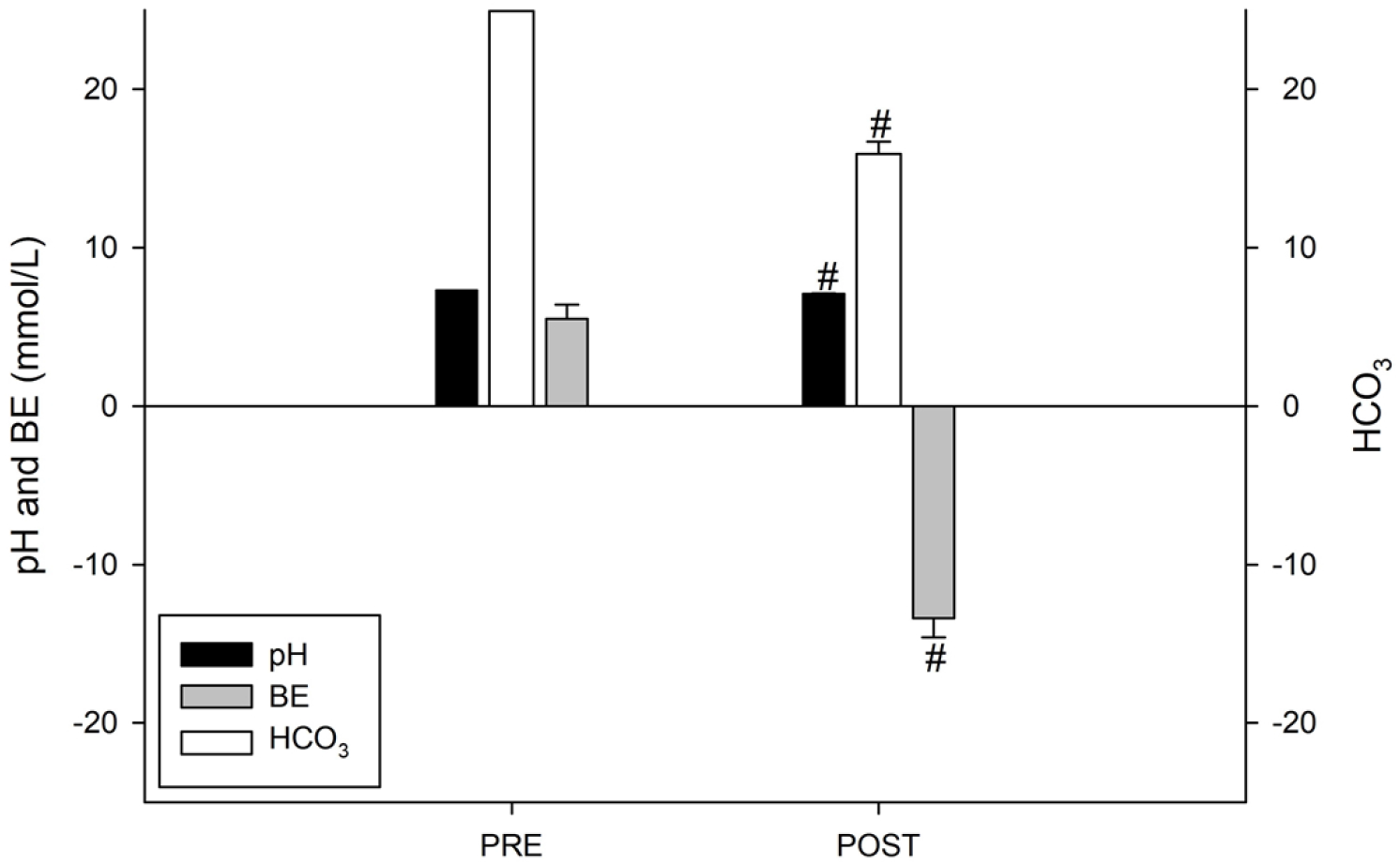
Exercise-induced acid-base balance responses.

Figure 3 presents a possibility for individualized observation of the sample participants, showing that the smaller the percentage variation in IL-12, the greater the phase to reach the anaerobic threshold in Km/h and the time to reach this same threshold, and the opposite is also true, where the greater the percentage change in IL-12, the shorter the phases and times to reach the AT. The visual observations in this graph were confirmed by the *Spearman* test, where a correlation coefficient of −0.900 was identified between IL-12 and the time to reach AT (P=0.0833) and of −0.872 between IL-12 and the phase to reach the AT (P=0.0833). These findings indicate a promising future in this line of correlations between immunometabolism and exercise.

**Figure 3.**
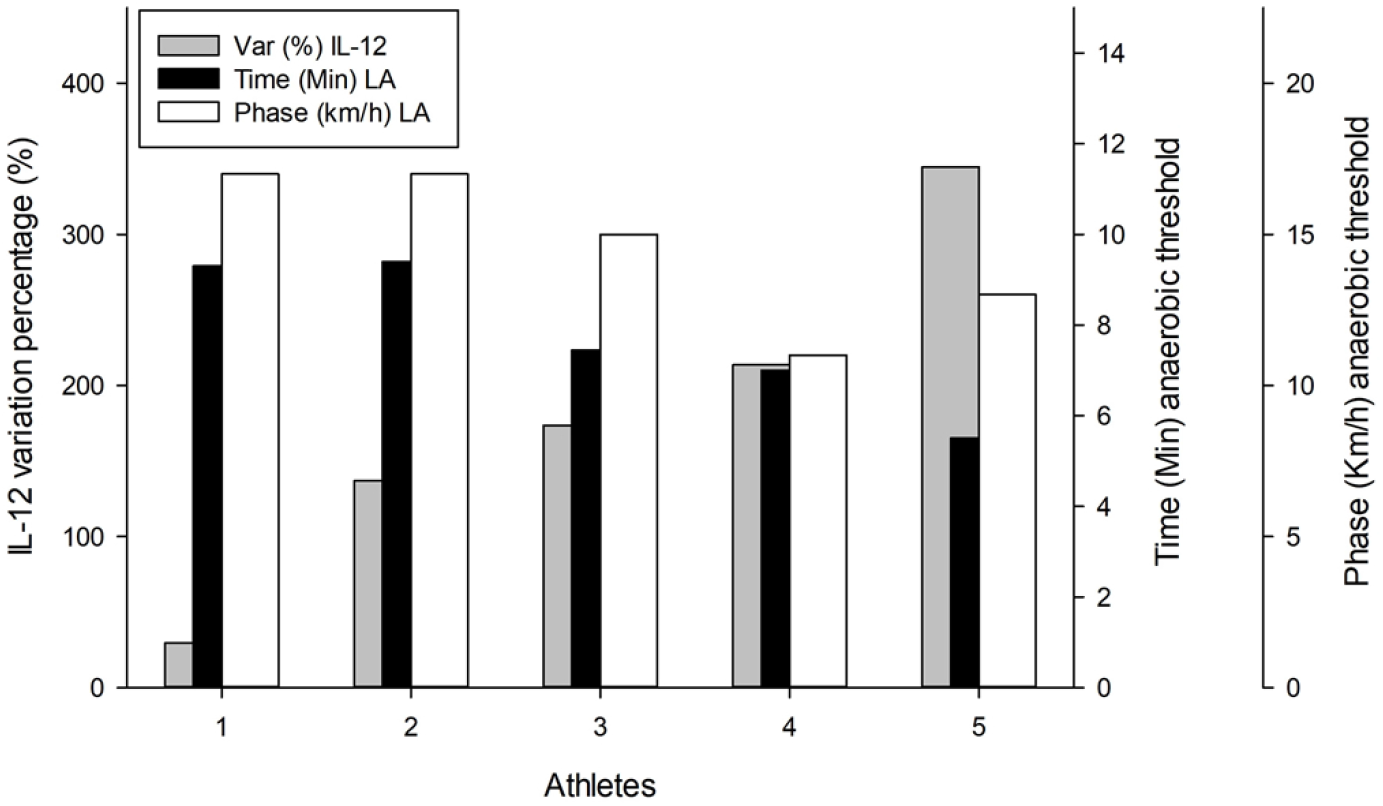
Individualized evaluation of the percentage variation in IL-12 and the time and phase to reach the anaerobic threshold.

The findings indicated in the previous paragraph demonstrate the need for an investigation of the correlation coefficients between all biomarkers and times in the present study. The values of the correlation coefficients obtained from the *Spearman* test are presented as a *heat map*, with a dark blue scale for coefficients from −1.0, passing through lighter shades of blue until reaching the color green for the zero value, and then passing through shades of yellow and light red until reaching dark red for the correlation of 1.0 (Figure 4).

**Figure 4.**
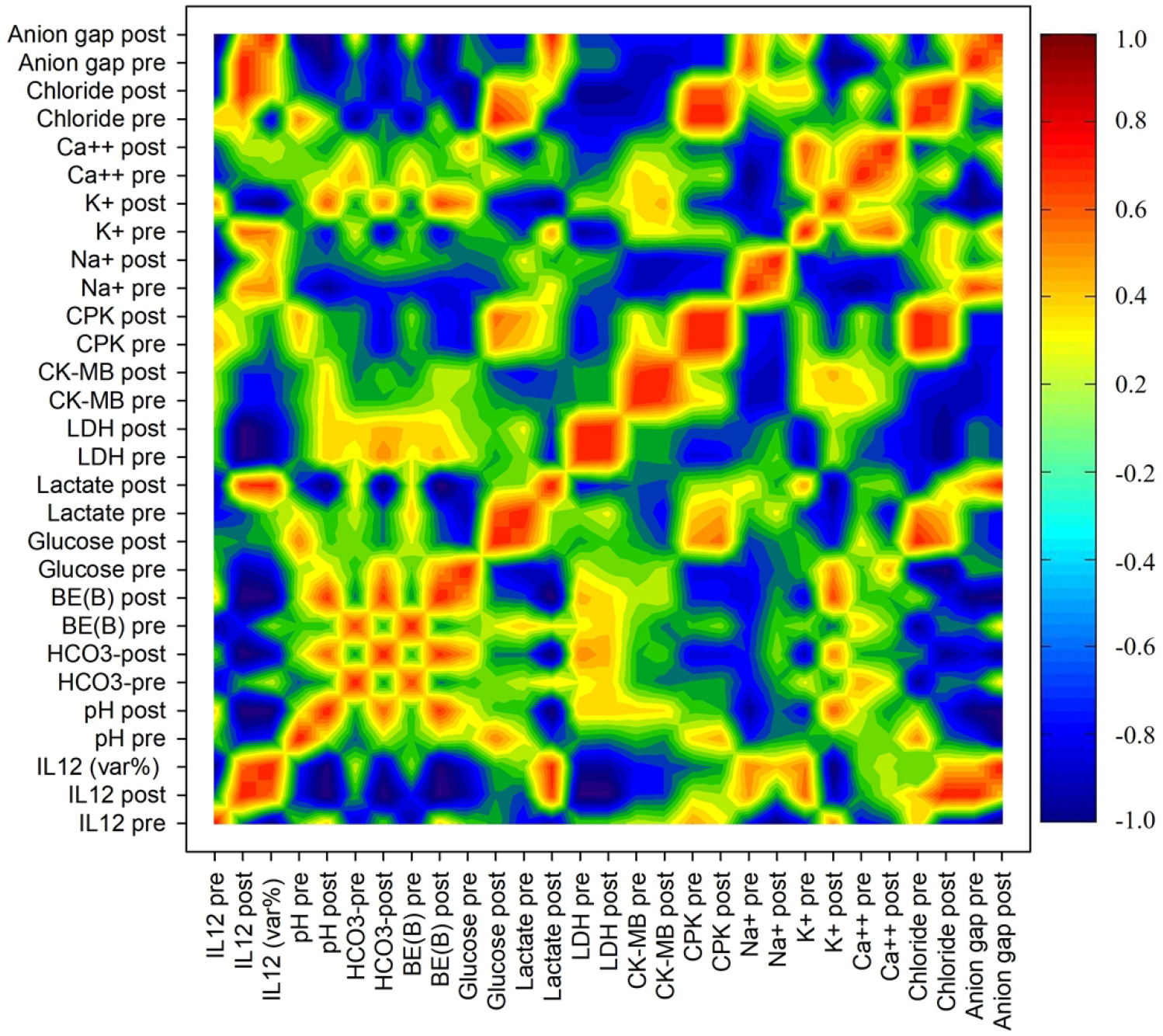
Heat map with data variation coefficients from the *Spearman* test.

Figure 4 presents intriguing positive correlations between post-exercise IL-12 and pre-anion gap (1.0 and P=0.333), post-exercise IL-12 and post chloride (1.0 and P=0.333), percentage change in IL-12 and post anion gap (1.0 and P=0.333) and percentage change in IL-12 and post lactate (0.943 and P=0.0167). In addition, negative correlations were observed between pre-IL-12 and post anion gap (−1.0 and P=0.333), post-IL-12 and pre-LDH (−0.943 and P=0.0167), post-IL-12 and post-LDH post (−0.943 and P=0.0167), post-IL-12 and post base excess (−9.943 and P=0.0167), post-IL-12 and post bicarbonate (−0.943 and P=0.0167), and post-IL-12 and post-pH (−0.943 and P=0.0167).

Figure 5, aims at a holistic and integrated analysis of the data, demonstrating a close network between acid-base balance biomarkers (Figure 5B) and great similarity between IL-12 and glycemia, LDH, and also acid-base balance (Figure 5B). All of the aforementioned findings will be discussed below.

**Figure 5.**
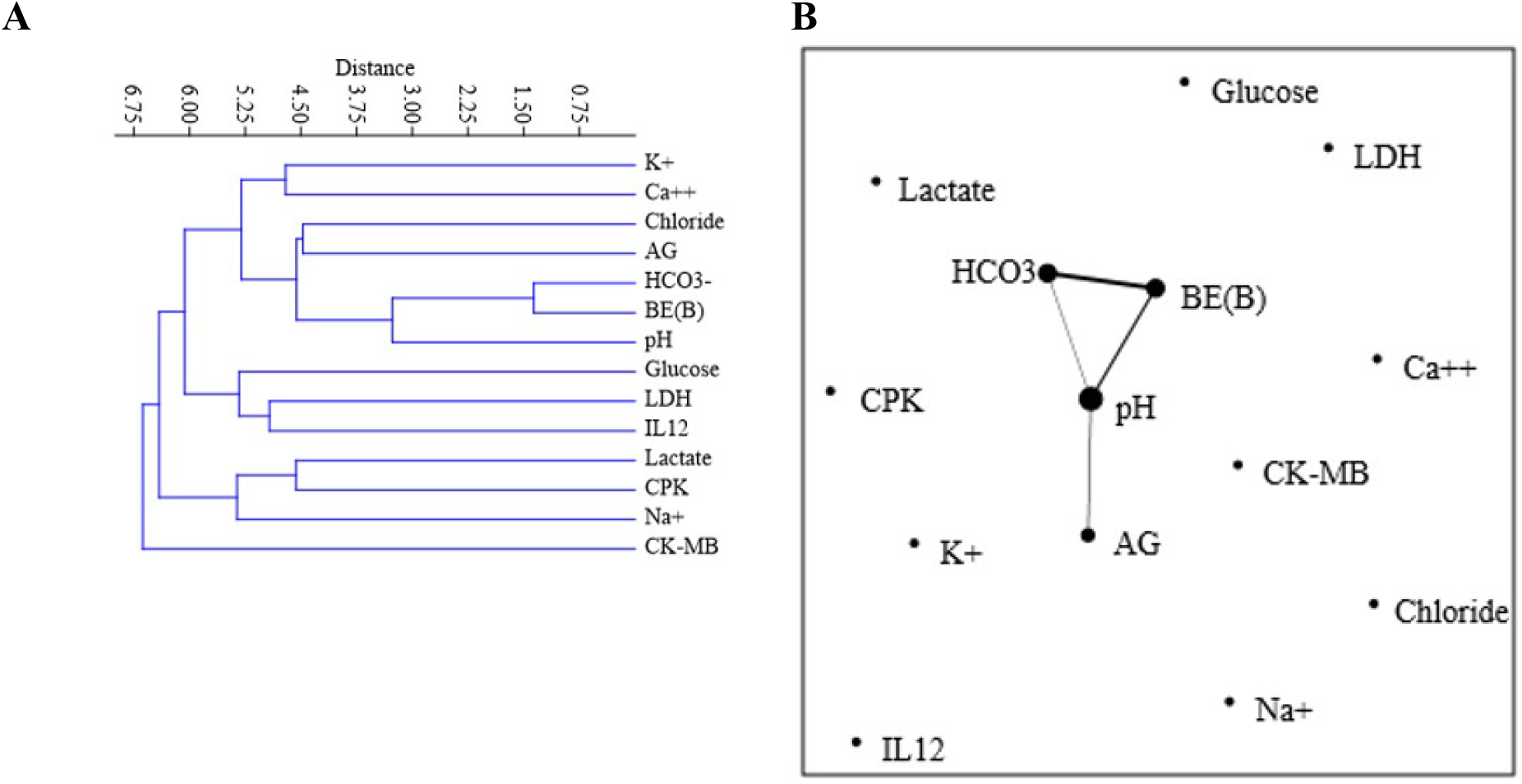
Dendrogram (A) and network plot with Fruchterman-Reingold algorithm (B) after calculating the Euclidean Similarity index.

## Discussion

The scarcity of sports studies involving immunometabolism makes more accurate analysis difficult, which is easily understood since this type of analysis is still considered embryonic, leaving many questions unanswered.

Nutrients absorbed from the diet can be divided into macronutrients and micronutrients, the first group being the most abundant, larger in size and able to donate energy for metabolism, and the second being smaller in size and quantity and with adjuvant functions in the functioning of the immunometabolism [28–30]. From the energy point of view of rapid use, carbohydrates stand out, presented as monosaccharides (aldoses or ketoses), disaccharides, or polysaccharides. The disaccharides include sucrose (glucose + fructose), lactose (glucose + galactose), and maltose (glucose + glucose), with glucose present in all [31–33].

Glycemic control represents an important factor for homeostasis, with important actors in this control, including hypoglycemic hormones, such as insulin, and hyperglycemic hormones, such as glucagon that acts on the hepatic tissue and epinephrine on the muscle, since these are the tissues that store glucose in the form of glycogen [34–37].

Increased blood glucose signals the pancreas to increase insulin secretion, which, when binding to enzyme-linked receptors contained in the membrane of different tissue families, promotes the opening of glucose transporters, with glucose uptake by cells [38–40].

Upon entering cells, glucose is rapidly converted to glucose-6-phosphate to promote glucose trapping and prevent it from leaving through a new channel opening [41–43]. From there, the energy status will determine whether it will be reserved in the form of glycogen (liver and muscle cells) or follow a multi-step metabolic pathway called glycolysis, aimed at the formation of adenosine triphosphate, which is the main energy currency of metabolism [44–46].

When metabolized in this way, pyruvate will be formed, which can be used as a substrate for the pyruvate dehydrogenase complex, forming acetyl-coenzyme A, which can be oxidized in the Krebs cycle, or follow other paths such as lipogenesis, also dependent on of energy status [46–48], or used as a substrate for enzyme lactate dehydrogenase (LDH), receiving two protons from the medium and forming lactate for cellular Ph homeostasis [49–51].

During exercise there is an increase in energy demand, with degradation of glucose to form energy, this demand promotes accumulation of protons in the intracellular environment, requiring greater formation of dose-dependent lactate (intensity or duration dependent) that will be released in the blood [52–54].

This lactate formed and available in the blood will be converted again into glucose in the liver and return to the blood, this conversion being through a metabolic pathway known as the Cori cycle [28,55,56].

In the present study, the running test on a treadmill promoted an average increase in glycemia of 58% and in lactatemia of 607% in only 13 minutes of testing. It is expected by the literature that, in general, approximately 15% of the appearance of glucose in the plasma is due to the activity of the Cori cycle [56], but this formation can be increased in highly trained individuals and during exercise [57], which could explain the 50% increase found in the studied sample.

Also very important for homeostasis, the acid-base balance of body tissues and fluids, especially blood, has intrigued physiologists and clinicians for some time [58]. The behavior of acids, bases, and salts in aqueous solutions is due to the activities of their ions, that is, if the hydrogen ions (protons) are in excess in relation to the hydroxyl ions, a solution is acidic, while if the opposite occurs, the solution is alkaline [59].

The buffer system made up of bicarbonate (HCO3-) and carbonic acid (H2CO3), essential in this balance, has special characteristics in body fluids, with the kidneys being responsible for their absorption and excretion in order to achieve balance [60].

Until 1948, when Singer and Hastings published their work describing the concept of a buffer base, acid-base problems were interpreted within the Henderson-Hasselbalch equation that describes the relationship between pH, pCO (or H2C03), and bicarbonate. The buffer base added a new parameter to separate airway and non-respiratory i.e., metabolic alterations [61].

Clinical assessment of acid-base imbalance relies on measurements taken in the blood, as part of the extracellular compartment. However, much of the metabolic importance of these disorders concerns intracellular events [62].

The kidneys have the predominant role in regulating systemic bicarbonate concentration and therefore the metabolic component of acid-base balance. This function of the kidneys has two components: reabsorption of virtually all filtered HCO3(−) and production of new bicarbonate to replace that consumed by normal or pathological acids [63]. However, not only bicarbonate enters this equation, the concentration of electrolytes is also important in this balance [64]. The anion gap equation is calculated from the sum of the serum concentrations of chloride and bicarbonate subtracted from the serum concentration of sodium [65]. Thus, lower serum bicarbonate levels and higher anion gap levels have been associated with lower cardiorespiratory fitness in adults [66].

Bicarbonate, however, is not the only base of the human organism. The plasma buffer base, which is nearly equal to the sum of bicarbonate and albumin anions, may be increased by base excess or by increased albumin concentration [67].

The present study protocol induced a 3% reduction in blood pH, a 48% reduction in bicarbonate, and a 529% reduction in base excess, with an elevation of the anion gap of 232%. As previously stated, this appears to be due to the decreased ability of acid-base homeostasis by respiration, promoting consumption of bases to provide balance and prevent metabolic acidosis.

Regarding performance and metabolism, the concept of anaerobic threshold (AT), introduced during the 1960s, can be detected by several methods suggested as new ‘thresholds’ based on the variable used for its detection, i.e., lactate, ventilatory threshold, glucose threshold [68]. Several tests and protocols have been created as non-invasive parameters for determining this threshold from exercise tests [69].

In the present study, when evaluated as an average, the anaerobic threshold was reached in 7:52 minutes, in the 14.9 Km/h phase, with a heart rate of 163 beats per minute, absolute power of 524 W, and absolute VO2 of 3.12 l.min. When carrying out an individual assessment of this threshold in the five athletes with an IL-12 dosage, it was possible to observe from the Spearman test a correlation coefficient of −0.900 between IL-12 and the time to reach the AT and of −0.872 between IL-12 and the phase to reach the AT. This negative correlation is made clear in figure 8, where it is possible to identify an inverse correlation between the IL-12 concentration and the Anaerobic Threshold, suggesting a relationship between this cytokine and performance.

Also noteworthy is the negative correlation observed in figure 9 between IL-12 and the enzyme LDH (−0.943) and the positive correlation between this same cytokine and lactate (0.943). These data provide robustness to the hypothesis raised in the above paragraph. In this same figure, it is also possible to observe a negative correlation between IL-12 and the anion gap (−1.0), and base excess (−0.943), bicarbonate concentration (−0.943), and pH (−0.943), in addition to a positive correlation between IL-12 and chlorides (1.0). These data also suggest a relationship between this cytokine and acid-base balance, but this relationship needs to be carefully evaluated.

## Conclusion

The present sportomic study on immunometabolism, in addition to presenting the stress generated by the treadmill test performed in high-level running athletes, suggested a possible correlation between interleukin 12 (IL-12) and performance, metabolism, and acid-base blood balance, without stating whether this cytokine modulates or is modulated by the aforementioned performance or balance. This investigation should be further developed in future studies.

It is also suggested that the removal of lactate and reestablishment of glycemia by the CORI cycle, with the expectation that around 15% of glycemia is formed by this cycle, may exceed this estimate in trained athletes, with a better metabolism performance according to the level of training.

## Declaration of Data Availability

All data generated or analyzed during this study are included in this published article (and its accompanying information files).

## Acknowledgements

We thank the athletes who are members of the athletics association of the city of Barra do Garças – MT, Brazil for agreeing to participate in the study, and especially to the Institute of Cardiology and the Chrono-immunomodulation laboratory, at the Federal University of Mato Grosso (UFMT), Araguaia Campus, Mato Grosso, Brazil.

## Authors’ contributions

IVC, LCOG, SGLR, and AMMN conceived and designed the study. IVC, LCOG, SGLR, and AMMN collected the data. IVC, LCOG, SGLR, JSSL, ELF, AHF, and AMMN analyzed the data and wrote the manuscript. All authors read and provided critical feedback on the manuscript prior to approval.

## Additional information

The authors declare that there are no conflicting interests.

---

**Supplementary table 1.** Data referring to the exercise test and cardiovascular aspects measured.

|  | **Mean** | **Median** | **SD** | **SE** |
| --- | --- | --- | --- | --- |
| **MVV (L/min)** | 163.3 | 162.2 | 13.7 | 4.3 |
| **FEV1 (L/min)** | 4.4 | 4.3 | 0.4 | 0.1 |
| **RD (Sec)** | 831.3 | 829.5 | 51.9 | 16.4 |
| **MHR1 (bpm)** | 198.8 | 200.5 | 5.5 | 1.7 |
| **MHR2 (bpm)** | 191.5 | 193.0 | 10.0 | 3.2 |
| **%MHR (%)** | 96.4 | 96.2 | 5.2 | 1.6 |
| **VO<sub>2</sub>Max1 (mL/kg·min)** | 53.6 | 54.7 | 3.4 | 1.1 |
| **VO<sub>2</sub>Max2 (mL/kg·min)** | 75.2 | 75.6 | 3.9 | 1.2 |
| **% VO<sub>2</sub>Max (%)** | 140.8 | 140.1 | 11.4 | 3.6 |
| **TD (mi)** | 1.8 | 1.8 | 0.1 | 0.0 |
| **TD (Km)** | 2.9 | 3.0 | 0.2 | 0.1 |
| **VO<sub>2</sub>/HR1 (ml/b)** | 16.3 | 16.1 | 1.5 | 0.5 |
| **VO<sub>2</sub>/HR2 (ml/b)** | 23.9 | 23.9 | 2.9 | 0.9 |
| **% VO<sub>2</sub>/HR2 (%)** | 146.4 | 148.4 | 13.2 | 4.2 |
| **MEV1 (BTPS) (L/MIN)** | 163.3 | 162.2 | 13.7 | 4.3 |
| **MEV2 (BTPS) (L/MIN)** | 162.0 | 156.1 | 14.6 | 4.6 |
| **% MEV (%)** | 99.4 | 100.6 | 8.0 | 2.5 |

MVV – Maximal Voluntary Ventilation; FEV1 – Forced Expiratory Volume; RD – Duration of the Test; MHR – Maximum Heart Rate (beats per minute); VO<sub>2</sub>Max – Maximum Rate of Oxygen Consumption Measured During Incremental Exercise; TD – Distance Traveled (Miles and Km); MEV – Maximum Expiratory Volume; (1) Estimated value (2) Measured value; (%) represents the percentage rate between redacted and measured values.

**Supplementary table 2.** Exercise-induced alterations in key biomarkers – mean (SE).

|  | **PRE** | **POST** | **Percentage $\Delta$** |
| --- | --- | --- | --- |
| **Hematocrit (%)** | 42 (1.0) | 44 (1.1) | 4% |
| **pH\*** | 7.3 (0.01) | 7.1 (0.02) | -3% |
| **HCO<sub>3</sub>\* (mmol/L)** | 31.0 (0.83) | 15.0 (0.83) | -48% |
| **BE\* (mmol/L)** | 3.9 (0.74) | -13.4 (1.19) | -529% |
| **Glucose\* (mmol/L)** | 78.6 (2.93) | 121.9 (6.37) | 58% |
| **Lactate\* (mmol/L)** | 1.9 (0.14) | 13.0 (1.01) | 607% |
| **LDH (U/L)** | 443.7 (35.4) | 500.7 (35.9) | 14% |
| **Chloride (mmol/L)** | 102.7 (0.94) | 105.3 (1.20) | 2.6% |
| **Sodium \* (mmol/L)** | 140.5 (0.44) | 143.7 (0.67) | 2.2% |
| **Potassium (mmol/L)** | 3.9 (0.09) | 3.2 (0.06) | -16% |
| **Calcium (mmol/L)** | 1.0 (0.04) | 1.0 (0.04) | -0.6% |
| **Anion gap** | 6.7 (0.90) | 22.3 (1.35) | 232% |
| **IL-12 (pg/ml)\*** | 3.4 (0.5) | 7.8 (0.5) | 160% |

621 \* Significant statistical difference.

## Notes

### Competing Interest Statement

The authors have declared no competing interest.

